# The landscape of molecular chaperones across human tissues reveals a layered architecture of core and variable chaperones

**DOI:** 10.1101/2020.03.04.976720

**Authors:** Netta Shemesh, Juman Jubran, Mehtap Abu-Qarn, Eyal Simonovky, Omer Basha, Idan Hekselman, Shiran Dror, Ekaterina Vinogradov, Serena Carra, Anat Ben-Zvi, Esti Yeger-Lotem

**Author notes:** These authors contributed equally to this study (Netta Shemesh, Juman Jubran). Correspondence to: Prof. Esti Yeger-Lotem, Prof. Anat Ben-Zvi.

## Abstract

The sensitivity of the protein-folding environment to chaperone disruption can be highly tissue-specific. Yet, the organization of the chaperone system across physiological human tissues has received little attention. Here, we used human tissue RNA-sequencing profiles to analyze the expression and organization of chaperones across 29 main tissues. We found that relative to protein-coding genes, chaperones were significantly more ubiquitously and highly expressed across all tissues. Nevertheless, differential expression analysis revealed that most chaperones were up- or down-regulated in certain tissues, suggesting that they have tissue-specific roles. In agreement, chaperones that were upregulated in skeletal muscle were highly enriched in mouse myoblasts and in nematode’s muscle tissue, and overlapped significantly with chaperones that are causal for muscle diseases. We also identified a distinct subset of chaperones that formed a uniformly-expressed, cross-family core group conducting basic cellular functions that was significantly more essential for cell survival. Altogether, this suggests a layered architecture of chaperones across tissues that is composed of shared core elements that are complemented by variable elements which give rise to tissue-specific functions and sensitivities, thereby contributing to the tissue-specificity of protein misfolding diseases.

**Significance Statement:** Protein misfolding diseases, such as neurodegenerative disorders and myopathies, are often manifested in a specific tissue or even a specific cell type. Enigmatically, however, they are typically caused by mutations in widely expressed proteins. Here we focused on chaperones, the main and basic components of the protein-folding machinery of cells. Computational analyses of large scale tissue transcriptomes unveils that the chaperone system is composed of core essential elements that are uniformly expressed across tissues, and of variable elements that are differentially expressed in a tissue-specific manner. This organization allows each tissue to fit the quality control system to its specific requirements and illuminates the mechanisms that underlie a tissue’s susceptibility to protein-misfolding diseases.

## Introduction

Chaperones are highly conserved molecular machines that control cellular protein homeostasis (proteostasis). Across species, they promote *de novo* protein folding and protein maturation (1), protein translocation (2), protein-complexes assembly and disassembly (3), protein disaggregation and refolding (4), and protein degradation (5). In accordance with their fundamental roles, chaperones are abundant proteins. In human cell lines, for example, they were shown to compose ~10% of the total proteome mass (6).

Chaperones have been grouped into families based on their molecular mass, common domains, protein structure similarity, and common function (1). Families composing the main chaperone machinery modulate protein structure while not forming a part of the final protein complex, and include prefoldin (7), the small heat shock proteins (sHSP) (8), and the main ATP-hydrolyzing chaperones, HSP60 (9), HSP70 (10), HSP90 (11) and HSP100 (12). Families composing co-chaperones modulate the activity of main chaperones by regulating their ATPase cycle or the recognition, binding, or release of chaperone substrates, and include HSP10 (9), HSP40 (DNAJ) (13), nuclear exchange factors (NEFs) (14) and co-HSP90 (15). Folding enzymes that catalyze folding-accelerating reactions, such as peptidyl prolyl cis-tans isomerization or protein disulfide isomerization (16, 17), are also considered as chaperones.

The chaperone system is highly versatile. Most chaperones interact with multiple co-chaperones, and co-chaperones can interact with multiple chaperones, thereby modifying their function or the fate of their substrates. For example, the same HSP70 chaperone can interact with different HSP40 co-chaperones, altering its substrate specificity (13). Likewise, co-chaperones containing the TPR domain, such as Hop, interact with both HSP70 and HSP90, thereby targeting substrates for either folding or degradation, respectively (18). The versatility and robustness of the chaperone system is manifested in stress conditions, which lead to extensive upregulation of chaperones. However, the system also has limitations. Whereas chaperone overexpression typically improves the folding capacity of the chaperone system, overexpression of specific chaperones was shown to disrupt folding (19–22). For example, overexpression of the folding enzyme FKBP51 in a tau transgenic mouse model resulted in accumulation of tau and its toxic oligomers (19–22). Likewise, chaperone downregulation or genetic aberration can cause diseases, such as mutation in the NEF co-chaperone BAG3 that leads to myopathy (23). These observations imply that the quantitative composition of the chaperone system can improve or impair its proteostatic capacity.

The chaperone system has expanded considerably in evolution (24, 25). The HSP40 family, for example, expanded from 3 members in *E. coli* and 22 members in budding yeast to 49 members in human, whose distinct functionalities are not totally clear (13). Consequently, the chaperone system has been remodeled. While in prokaryotes the same chaperones carry both *de novo* protein folding and response to stress, unicellular eukaryotes evolved two separately-regulated chaperone systems, a basal system and a stress inducible system, each composing distinct members of the same chaperone families (26). This two-level organization is conserved in multi-cellular eukaryotes. Yet, multi-cellular eukaryotes are also composed of multiple cell types, tissues and organs, each having different proteomes and thus potentially different folding demands. For example, HSPB1 is required for actin and myofibril assembly and its depletion impairs cardiac progenitor fusion and heart tube formation (27).

Recent years were marked by multiple large-scale mappings of the proteomic and transcriptomic landscapes of tens of human tissues. Major efforts included the Human Protein Atlas (28), Fantom5 (29), and the Genotype Tissue Expression (GTEx) consortium (30). These resources enabled unprecedented quantitative views into the genes and proteins that make up physiological human tissues. These studies and others have revealed tissue-specific regulatory elements, molecular interaction networks, and functional mechanisms underlying traits and diseases (30–34). The chaperone system at large-scale was studied in the context of neurodegenerative diseases (35), heat stress (36) and cancer (37) via analysis of samples gathered from patients. These revealed different patterns of chaperone network deregulation. However, a systematic examination of the basal chaperone network in physiological human tissues has been lacking.

Here we harnessed transcriptomic profiles gathered by the GTEx consortium to systematically examine the basal chaperone system in various human tissues. We focused on 194 manually-curated chaperones, co-chaperones and folding enzymes. We found that in accordance with the fundamental role of the chaperone system, chaperones are significantly more ubiquitously and highly expressed across all tissues relative to other protein-coding genes, and are also more essential for growth. Nevertheless, differential analysis of chaperones expression across tissues showed that most chaperones have tissue-specific behaviors. A proteomic screen of mouse myocytes and a computational analysis of disease-causing chaperones showed that these behaviors tend to be conserved and functional. We further highlight a core set of chaperones that is uniformly expressed across tissues, and show them to be more essential for growth than other chaperones. This core subsystem still establishes tissue-specific functional networks, which are enhanced by their tissue-specific relationships with other chaperones. We propose that the combination of core and non-core chaperones constitutes a third-level organization of the chaperone system that is capable of supporting the different proteostatic demands of multi-cellular eukaryotes, and that this organization illuminates the phenotypic outcomes of chaperones aberrations.

## Results

### Chaperones are ubiquitously and highly expressed across human tissues

We manually curated a list of multi-substrate chaperones and co-chaperones of the main eukaryotic chaperone families, which drive basic cellular processes needed in living cells. These included sHSP, HSP40, HSP60/HSP10, HSP70, HSP90, prefoldin, and folding enzymes, as well as chaperones not structurally or functionally associated with a specific chaperone family, such as ER chaperones, which we denoted as “other” (Table S1). To analyze their expression across tissues, we used RNA-sequencing (RNA-seq) profiles that were made available by the GTEx consortium (30). We focused on tissues with at least 5 available profiles sampled from donors with traumatic injury fitting with sudden death, and avoided other death causes to limit the effect of treatment and stress on chaperone expression. This resulted in a total of 29 tissues, 488 profiles, and 194 expressed chaperones, co-chaperones and folding enzymes, henceforth referred to as chaperones (Table S1).

We first compared the presence of chaperones across tissues to that of other protein-coding genes. We considered a gene as present in a tissue if its median expression level in samples of that tissue was above a certain threshold (see Methods). We found that chaperones were significantly more ubiquitous than other protein-coding genes (p=3.9E-15, Kolmogorov-Smirnov test, Fig. 1A). For example, 74% of the chaperones were present in all tissues, relative to 47% of the protein-coding genes. Similar results were observed with other thresholds (Fig. S1). Next, we examined the expression levels of chaperones relative to protein-coding genes. Within each of the 29 tissues that we analyzed, the expression levels of chaperones were significantly higher (p<1.7E-7, Mann-Whitney, Fig. 1B), also when considering other thresholds (Fig. S1B).

**Figure 1.**
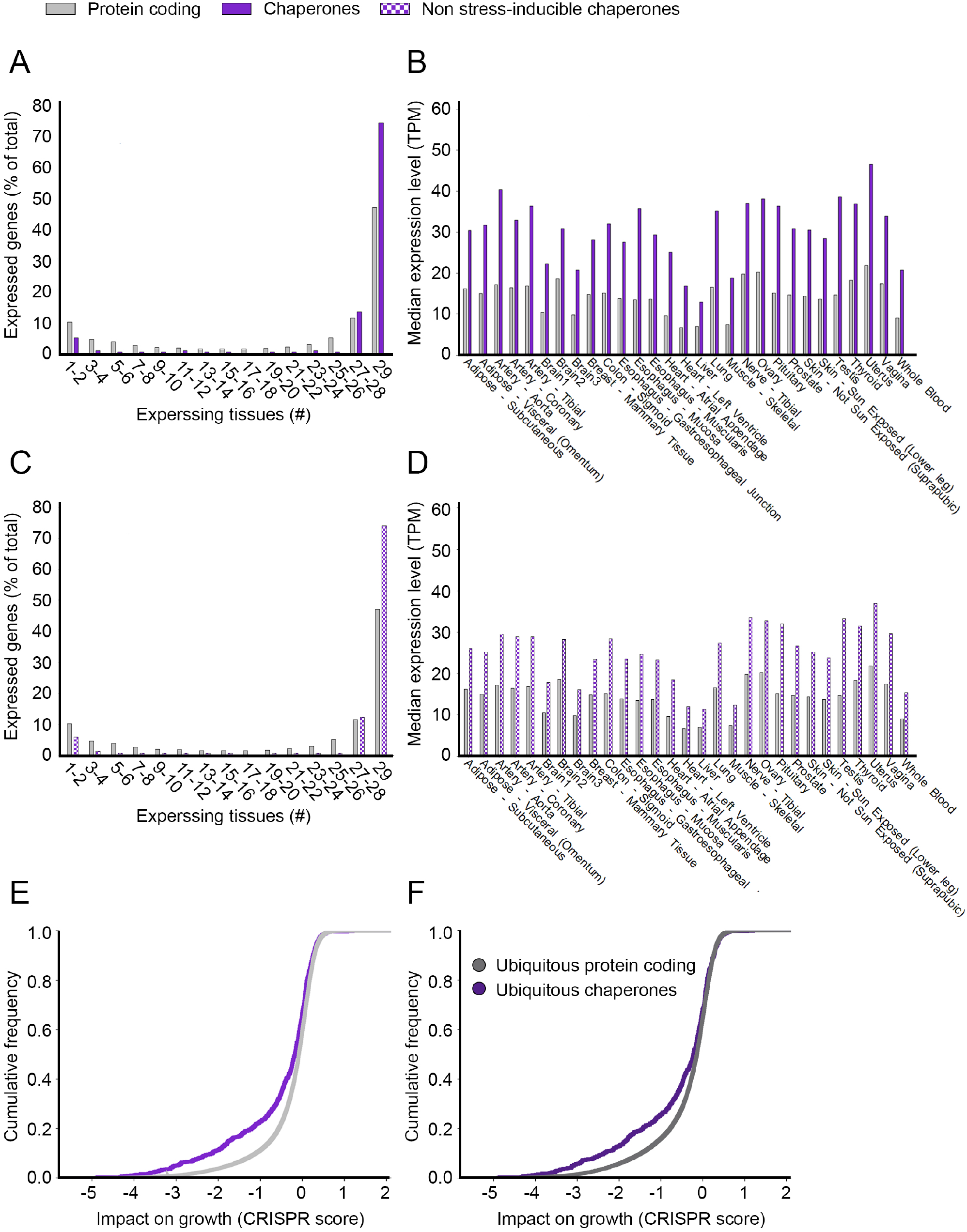
Chaperones are fundamental components of human tissues. A. The distribution of 194 chaperones and 17,689 other protein-coding genes by the number of tissues expressing them at a level of 1TPM or above. Chaperones are significantly more ubiquitously expressed than other protein-coding genes (p=3.9E-15, Kolmogorov-Smirnov test). B. The median expression levels per tissue of chaperones and other protein-coding genes. Only genes expressed at a level of 1TPM or above were considered. Chaperones tend to be significantly more highly-expressed across all 29 tissues (p<1.7E-7, Mann Whitney test). C. The distribution of 154 non-inducible chaperones and 17,689 other protein-coding genes by the number of tissues expressing them at a level of 1TPM or above. Non-inducible chaperones are significantly more ubiquitously expressed than other protein-coding genes (p=6.7E-17, Kolmogorov-Smirnov test). D. The median expression levels per tissue of non-inducible chaperones and other protein-coding genes. Only genes expressed at a level of 1TPM or above were considered. Non-inducible chaperones tend to be significantly more highly-expressed across all 29 tissues (p<4.1E-5, Mann Whitney test). E. The cumulative distribution of the essentiality scores of 191 chaperones and 17,973 other protein-coding genes, as measured by the impact of their inactivation by CRISPR on cell growth (CRISPR score). Chaperones were significantly more essential than other protein-coding genes (p=8.5E-12, Kolmogorov-Smirnov test). F. The cumulative distribution of the essentiality scores of the subsets of 165 chaperones and 10,324 other protein-coding genes that were expressed ubiquitously in all tissues. Chaperones tend to be more essential even when compared to other ubiquitously expressed genes (p=9.2E-7, Kolmogorov-Smirnov test).

Since stress-induced chaperones are known to be highly expressed in stressful conditions, they could be driving the high expression that we observed. To test for this, we defined a subset of 40 known stress-induced human chaperones, which were determined using a metanalysis of microarray data (36), and found them to be more highly expressed than non-inducible chaperones (Fig. S1C). However, even upon excluding them from the analysis, the remaining non-inducible chaperones were still significantly more ubiquitous (p=6.7E-17, Fig. 1C and Fig. S1D,) and more highly expressed than those of protein-coding genes (p<4.1E-5, Fig. 1D and Fig. S1E).

To further examine to what extent chaperones are fundamental components of living cells, we utilized large-scale data of gene essentiality measured by the impact of CRISPR-induced individual gene inactivation on growth rate of human lymphoma and myeloid leukemia cell lines (38). We found that relative to inactivation of other protein-coding genes, inactivated chaperones lowered growth rates significantly, and were thus considered more essential for growth (p=8.5E-12, Kolmogorov-Smirnov test, Fig. 1E). The higher essentiality of chaperones could be common to other ubiquitously expressed protein-coding genes. However, upon limiting our analysis to the subsets of ubiquitously expressed protein-coding genes and chaperones, chaperones were still more essential for growth (p=9.2*E-7, Kolmogorov-Smirnov test, Fig. 1F). Altogether, chaperones appear to constitute a ubiquitous and highly expressed cellular system that is fundamental for cell growth.

### Chaperone expression levels vary across human tissues

Despite the ‘house-keeping’ functionality of chaperones, different tissues have different folding, assembly, and maintenance demands. For example, the demands of the sarcomere in muscle cells, which contain titin, the largest protein in the human body (39, 40) likely vary from those of synaptic neurotransmission in neurons, where the membrane protein synaptic vesicle protein 2 (SV2) is amongst the most abundant and conserved components (41, 42). Such demands become phenotypically apparent in face of mutations. To test whether these demands are met by tissue-specific chaperones, we gathered from the literature 62 tissue-selective heritable disorders known to be caused by aberrations in 43 distinct chaperones, and analyzed the expression patterns of these chaperones across tissues (see Methods, Table S2). In contrast to the tissue-specific clinical manifestation of the chaperone-based disorders, most disease-causing chaperones were expressed ubiquitously across tissues (Fig. 2A), also when considering other expression thresholds (Fig. S2). Thus, mere presence of a chaperone in a tissue is not sufficient for meeting the tissue-specific folding demands.

**Figure 2.**
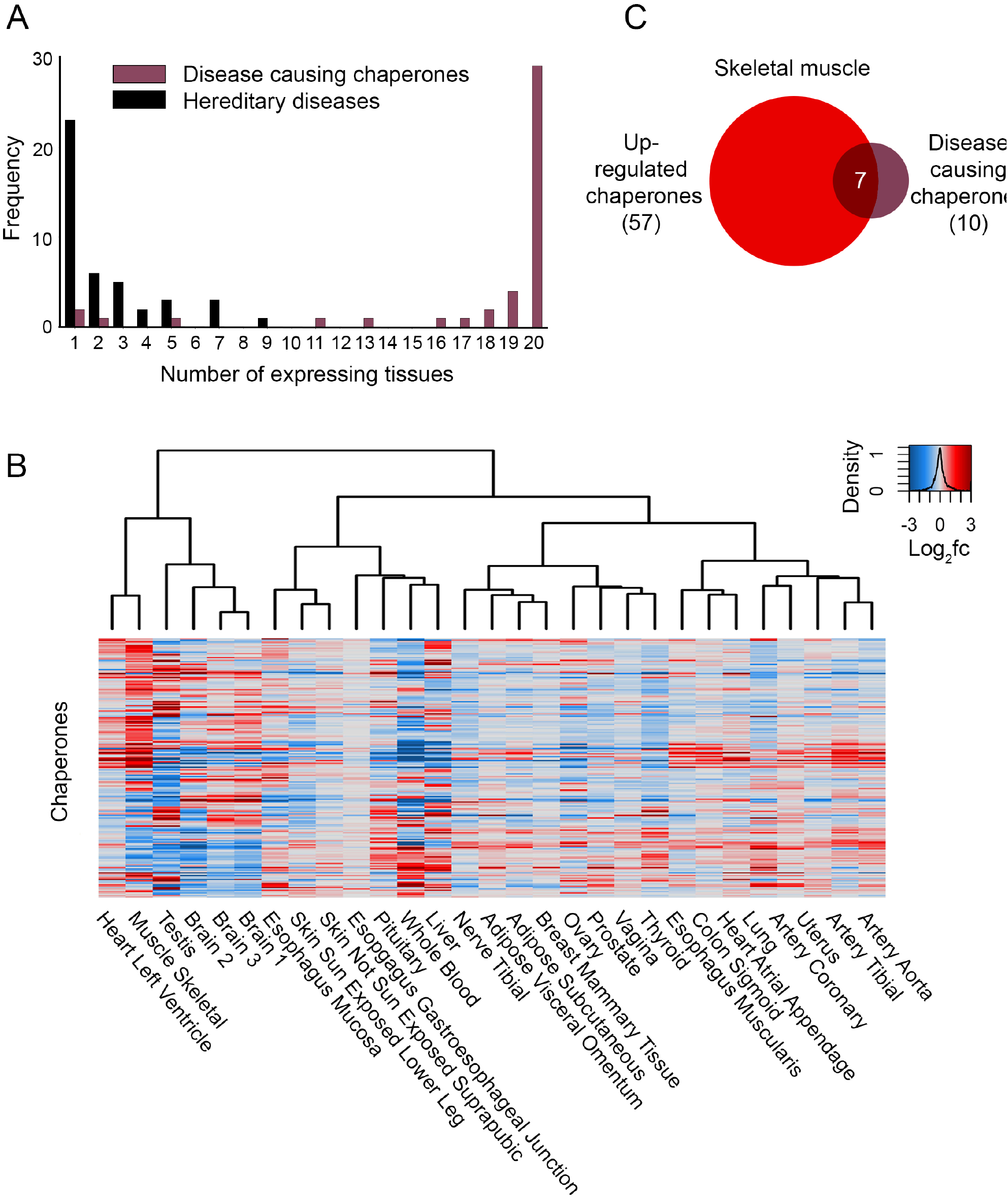
The variable expression of chaperones across human tissues. A. The distribution of 43 chaperones that are causal for 62 Mendelian diseases by the number of tissues expressing them at a level of 1TPM or above, and the number of tissues in which their diseases manifest clinically. We united sub-parts of the same tissue (e.g., adipose subcutaneous and adipose visceral omentum were united into a single adipose tissue), ending up with 20 united tissues. In contrast to the ubiquitous expression of chaperones across tissues, most diseases are highly tissue-specific, also when considering other expression thresholds (Fig. S2). B. A clustered heatmap showing the differential expression of 194 chaperones across tissues. Differential expression was computed for each chaperone per tissue relative to all other tissues. The log_2_ fold-change values (log_2_FC) are shown; red and blue denote upregulated and downregulated chaperones, respectively. Physiologically-related tissues often clustered together. C. The overlap between 57 chaperones that were up-regulated in human skeletal muscle (log_2_FC ≥ 1) and the 10 chaperones that are causal for heritable muscle disorders (7 chaperones, p=0.0078, Fisher exact test).

Next, we asked whether the expression levels of chaperones tend to vary across tissues. For this, we computed per tissue the differential expression of chaperones in the given tissue relative to all other tissues (43). We then clustered chaperones and tissues hierarchically according to their differential expression profiles (Fig 2B). Notably, physiologically related tissues, such as skeletal muscle and cardiac muscle, or different brain tissues, clustered together, supporting the biological relevance of the differential expression patterns. Some chaperones did not vary by more than 2-fold in any tissue, such as HSPA8 of the HSP70 family and HSP90AA1 of the HSP90 family, and were thus considered to be uniformly expressed across tissues. However, most chaperones (162, 83.5%) showed a 2-fold change in at least one tissue, henceforth considered as variably expressed.

To examine the biological relevance of tissue-specific expression profiles, we asked whether tissue-specific changes in expression were related to tissue-specific phenotypes. We focused on muscle, since it showed a distinct differential expression profile relative to other tissues, and since our heritable disorders’ dataset contained many muscle-specific diseases. We found that chaperones that were at least 2-fold upregulated in muscle were enriched significantly for chaperones that are causal for muscle disease (p=0.0078, Fisher exact test, Fig. 2C). For example, DNAJB6 [log 2 fold-change (log2FC) of 2.24] and the NEF chaperone BAG3 (log2FC of 3.13) lead to Limb-gridle muscular dystrophy 1E (44), and myopathies, respectively (45–47). Thus, chaperones expression levels are likely designed to meet tissue-specific demands and consequently affect disease risks in a tissue-specific manner.

### A conserved chaperone expression pattern in muscle

Since our analyses relied on the GTEx transcriptomic dataset, which might not be informative of tissue proteomes, we decided to test chaperone expression patterns at the protein level. For this, we profiled the proteome of C2C12 mouse myoblast cell line before differentiation (day 0) and after 8 days of differentiation to myotubes (day 8, myotubes) or reserve cells (day 8, reserved cells; Fig. 3A). We then compared the differential proteomic profile of mouse muscle (day 8 myotubes vs day 0) to the differential transcriptomic profile of human skeletal muscle. The proteomic and transcriptomic profiles correlated significantly upon considering all protein-coding genes (r=0.65, p<2.2E-16, Spearman correlation, Fig. 3B) and only chaperones (r=0.48, p<4.4E-5, Spearman correlation, Fig. 3C). In contrast, no significant correlations were observed when we compared the differential proteomic profile of reserved cells to the differential transcriptomic profile of human skeletal muscle (Fig. 3D). These results support the functional relevance of the human muscle differential transcriptomic data.

**Figure 3.**
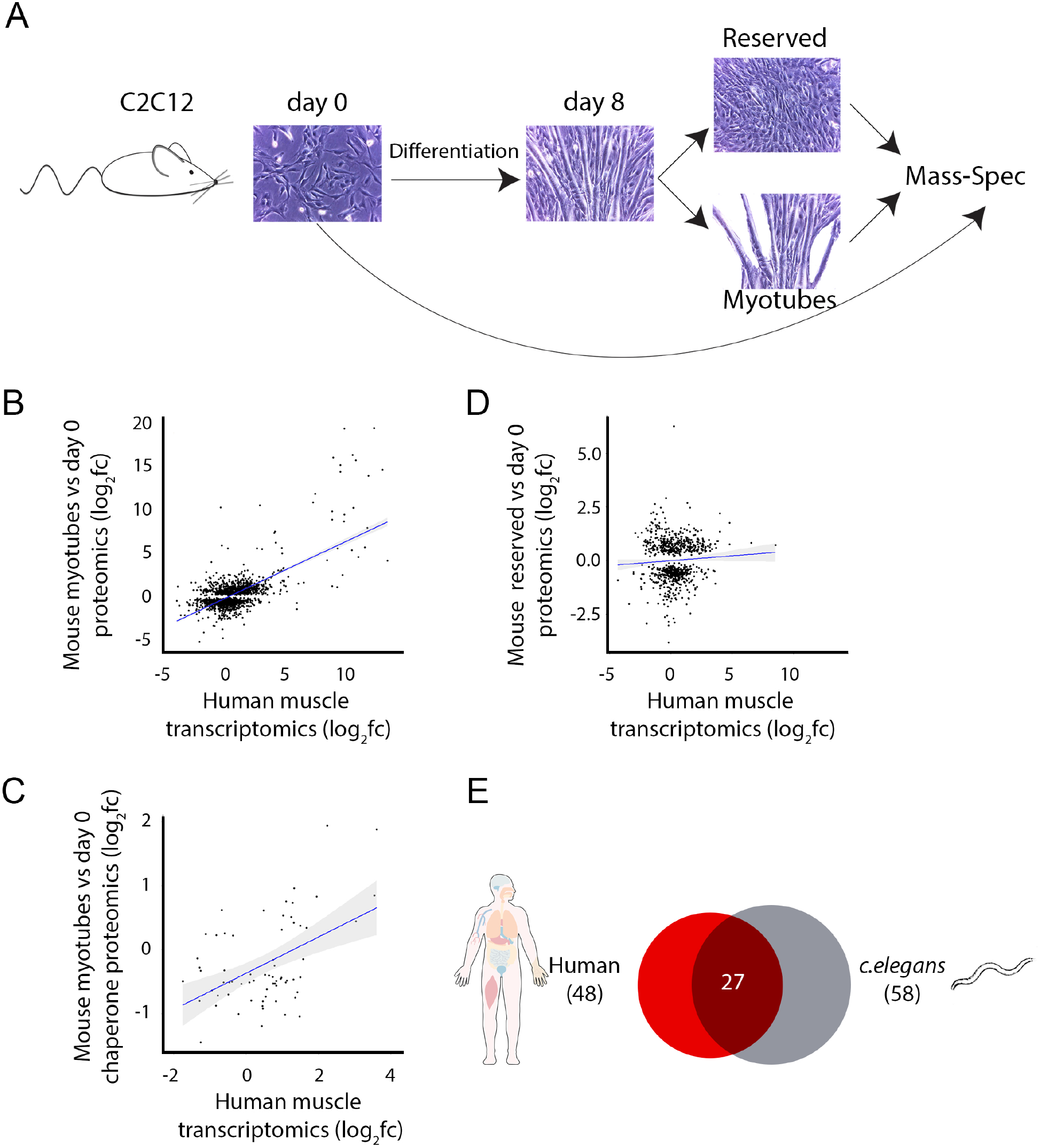
Chaperones function in muscle tissue is evolutionary conserved. A. The experimental pipeline. C2C12 mouse cells were grown to 95% confluency and differentiation was induced. After 8 days of differentiation, cells were separated to myotubes and reserved cells, and their proteomes were analyzed using mass spectrometry. B. The correlation between the differential proteomic profile of 1,605 proteins that were reliably measured in mouse myotubes versus undifferentiated cells, and the differential expression of their homologous genes in human skeletal muscle (r=0.65, p<2.2E-16, Spearman correlation). C. The correlation between the differential proteomic profile of 67 chaperones that were reliably measured in mouse myotubes versus undifferentiated cells, and the differential expression of their homologous genes in human skeletal muscle (r=0.48, p<4.4E-5, Spearman correlation). D. The differential proteomic profile of 776 proteins that were reliably measured in mouse reserved cells versus undifferentiated cells does not correlate with the differential expression of their homologous genes in human skeletal muscle (r=0.062, Spearman correlation). E. The overlap between 58 chaperones that were enriched in *C. elegans* muscle and 48 homologous chaperones that were upregulated in human skeletal muscle (p=0.0009, Fisher exact test). Only the 157 human chaperones with homologous genes in *C. elegans* were considered in the analysis.

We further tested the functional conservation of muscle chaperones by comparing between human and the evolutionary distant multicellular organism *Caenorhabditis elegans*. We used a set of *C. elegans* chaperones that were enriched in muscle (19), and compared them to orthologous chaperones that were upregulated in human muscle (see Methods). We found that the two subsets overlapped significantly (p=0.0009, Fisher exact test, Fig. 3E). For example, the HSP40 protein DNAJB6 was upregulated in human and its *C. elegans* homolog, *dnj-24*, was enriched in *C. elegans* muscle cells. Altogether, these observations suggest an inherent requirement for chaperones in muscle tissue function that is conserved in evolution.

### Chaperone families show variable behavior across tissues

Since members of each chaperone family share functional and structural attributes, we examined whether they also share expression patterns across tissues. We focused on their tendency for ubiquitous expression, for variable expression across tissues, and their essentiality for growth. To assess the ubiquitous expression of each family, we recorded the number of tissues in which chaperones were present (as in Fig. 1A). Almost all members of the ATP-hydrolyzing chaperone families,HSP60, HSP70 and HSP90, and the prefoldin family were ubiquitously expressed (present in >26 of the tissues). The only exception (1 of 40) was an HSP60 chaperonin that encodes a CCT subunit-like protein, CCT8L2, expressed specifically in testis. In contrast, while members of co-chaperone families were generally ubiquitously expressed, several of their members showed tissue-specific expression (Fig. 4A). These include five members of the HSP40 family, which are expressed in 1-2 tissues (at this threshold).

**Figure 4:**
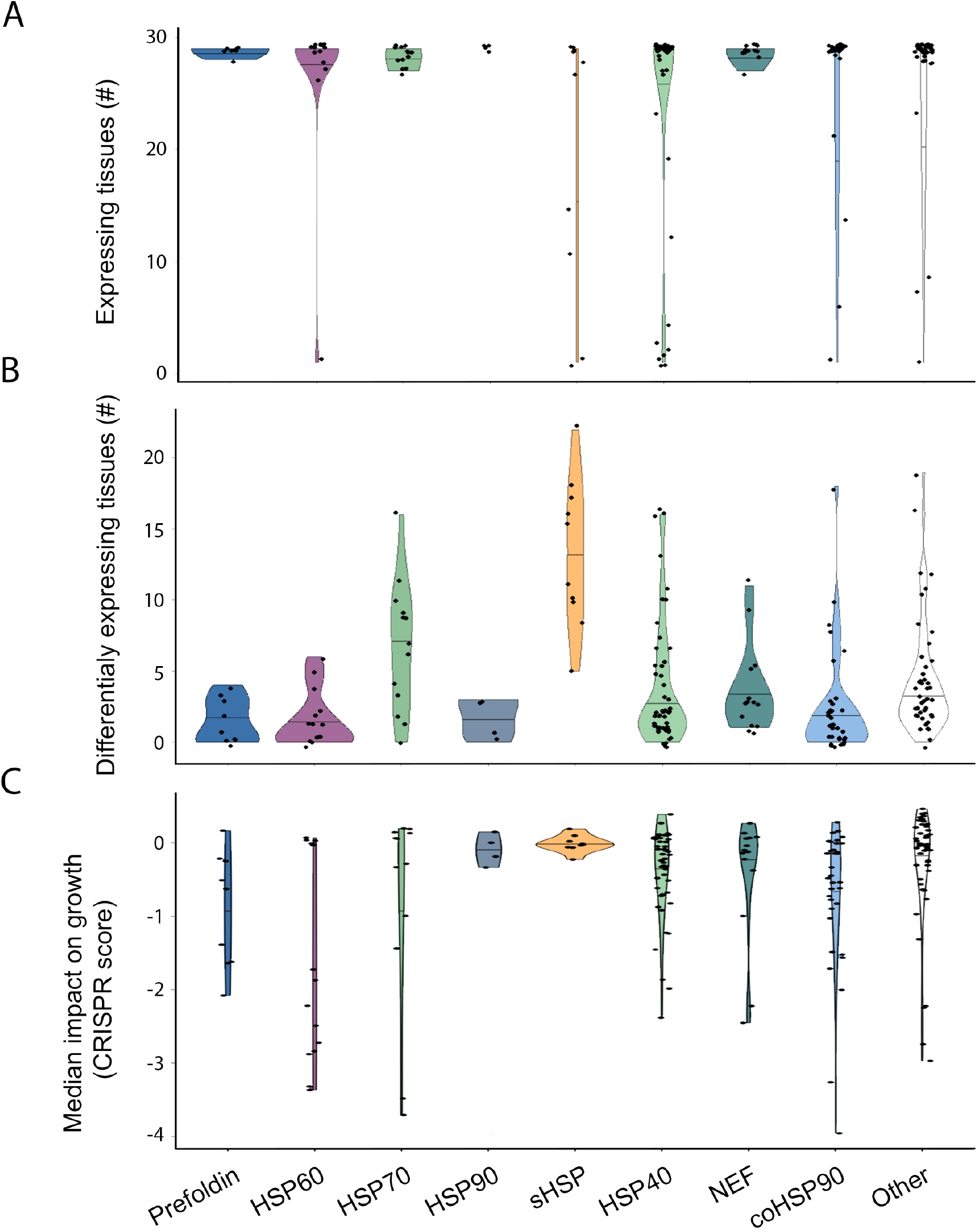
The heterogeneous expression and essentiality patterns of chaperone families. A. The distribution of the number of tissues expressing a chaperone, per family. A chaperone was considered as expressed in a tissue if its expression level was ≥ 1 TPM. Families encoding the main ATP-hydrolyzing chaperone machinery and the prefoldin family tend to be ubiquitously expressed across tissues. B. The distribution of the number of tissues in which a chaperone was differentially expressed (absolute (log_2_FC) > 1) per family. Except for HSP70, families encoding the main ATP-hydrolyzing chaperone machinery tend to be uniformly expressed across tissues. C. The distribution of the essentiality scores of chaperones, per family. For each chaperone its median essentiality score was considered. Chaperone essentiality was highly diverse in most chaperone families.

To assess the tendency of family members for uniform versus variable expression across tissues, we associated each chaperone with the number of tissues in which it was variably expressed (Fig. 4B). Families that were ubiquitously expressed, including prefoldin and two of the three families encoding the main ATP-hydrolyzing chaperone machinery, were also uniformly expressed across tissues, in agreement with their key functions in proteostasis regulation. The remaining family, HSP70, was variably expressed, but not its more essential members, including the main cytosolic member HSPA8, the ribosome associated member HSPA14, the ER member HSPA5 and the mitochondrial member HSPA9, which were more uniformly expressed (Fig. 4B). Likewise, most co-chaperone families had both uniform and variably expressed members. The only exception was the sHSP family, all members of the sHSP family were variably expressed across tissues (Fig. 4B).

Lastly, we assessed the essentiality for growth of each family (Fig. 4C). We found that chaperone essentiality was highly diverse in most chaperone families. For example, members of the HSP70 family that are required for basic cellular functions, such as HSPA8, HSPA14, HSPA5 and HSPA9, were highly essential, while stress inducible members, such as HSPA6 and HSPA1A, were not. Likewise, most CCT subunits were essential for growth, but not the testis-specific CCT8L2. In contrast, all the members of the sHSP family were not essential for growth, fitting with them having tissue-specific roles. Indeed, 5/10 of the sHSP chaperones are causal for tissue-specific diseases; for example, mutations in HSPB1, HSPB3, HSPB5 (CRYAB) and HSPB8 are causal for diseases of the neuro-muscular system (48–53), namely motor neuropathies and myopathies (Table S2). In summary, whereas most members of the HSP60, HSP90, and prefoldin families are ubiquitously and uniformly expressed and all members of the sHSP are variably expressed, making these features family-based, members of most chaperone families show distinct features that are likely associated with distinct functions across tissues.

### Core chaperones versus variable chaperones

We next decided to examine whether the small subset of 32 (16%) chaperones that were expressed uniformly across tissues, henceforth denoted as core chaperones, shared common functional features. Literature mining of the cellular processes in which these core chaperones participate showed that they covered a variety of basic processes, including: *de novo* folding and protein maturation and assembly; ER-associated targeting, transport, folding and degradation; and translocation into the mitochondria (Fig. 5A). Next, we asked whether they included known diseases-causing genes (see Methods). In contrast to variable chaperones, which were significantly enriched for disease-causing genes (p=0.04, Fisher exact test), only 3/32 core chaperones were disease-causing (p=0.99, Fig. 5B). This suggests that core chaperones do not cause diseases either because they are more functionally redundant with other chaperones, or, inversely, because their aberration is lethal. For example CCT5 CRISPR knockout (Cct5^em1(IMPC)Tcp^) mutant mice showed preweaning lethality. To quantitatively distinguish between the two alternatives, we compared between the impact on growth of mutations in core and variable chaperones (38). We found that core chaperones were significantly more essential for growth than variable chaperones (p=5.4E-15, Kolmogorov-Smirnov test, Fig 5C), also when limiting the variable chaperone to those that were expressed ubiquitously (p=2.7E-12). Lastly, we compared the median expression levels of core versus variable chaperones across tissues (Fig. 5D). Core chaperones were significantly more highly expressed than variable chaperones (p<2.2E-16, Mann-Whitney test). Taken together, we propose that core chaperones establish an essential subset of chaperones with stringent, uniform demand across tissues.

**Figure 5.**
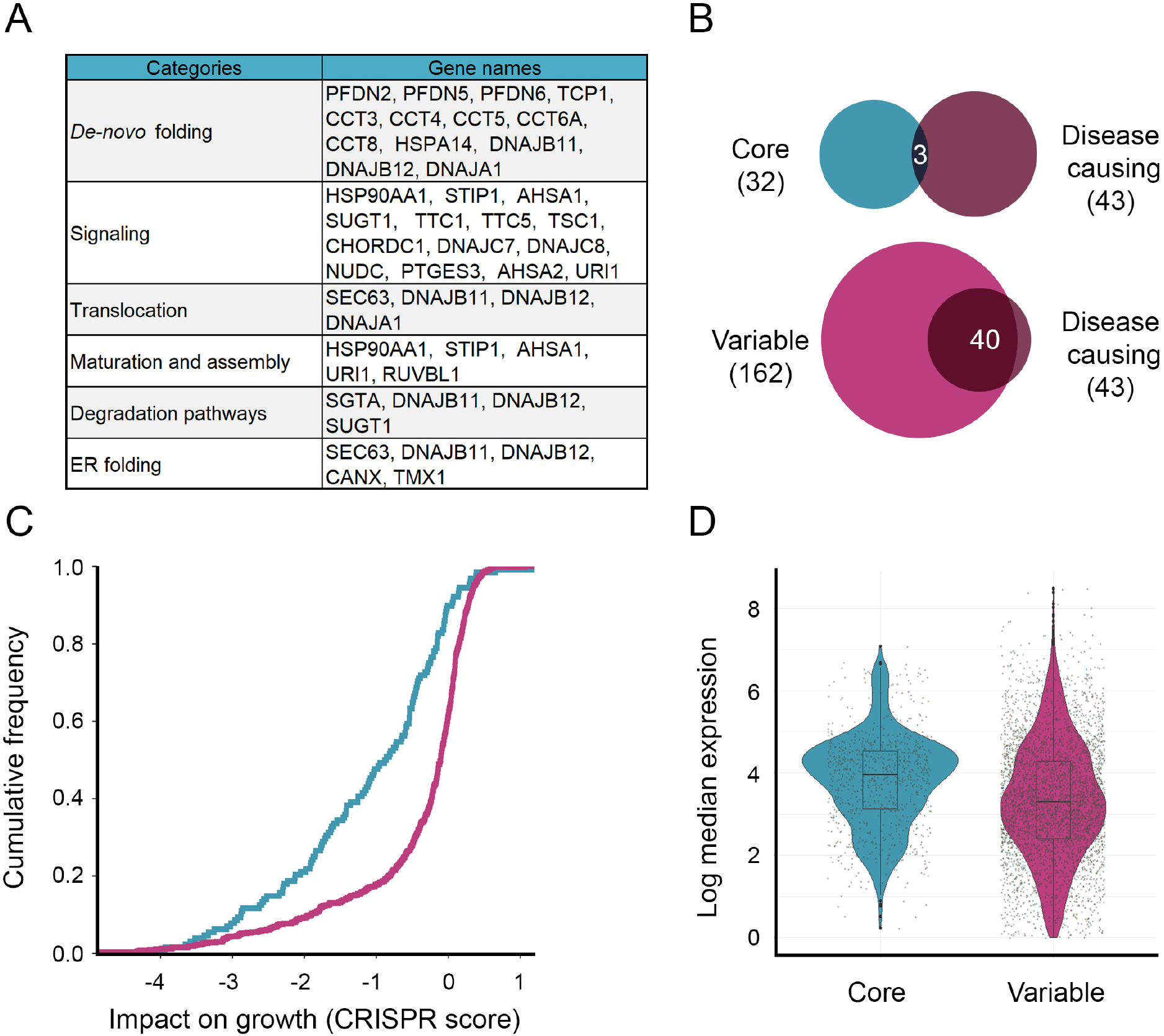
Core chaperones are uniformly expressed and required across tissues. A. The functional roles of the 32 core chaperones. B. The overlap between core chaperones (top) or variable chaperones (bottom), and the chaperones that are causal for Mendelian diseases. Core chaperones tend to be depleted of disease-causing chaperones, whereas variable chaperones tend to be enriched for them (p=0.04, Fisher exact test). C. The cumulative distribution of the essentiality scores of core versus variable chaperones, as measured by the impact of their inactivation by CRISPR on cell growth (CRISPR score). 32 core and 159 variable chaperones for which CRISPR scores were available were considered. Core chaperones are significantly more essential (p=5.4E-15, Kolmogorov-Smirnov test). D. The median expression levels of core versus variable chaperones. Core chaperones are significantly more highly expressed (p<2.2E-16, Mann Whitney test).

### Chaperone networks differ between tissues

Chaperones act as part of a functional network. Previous studies analyzed the physical interactions of a subset of chaperones in tissue culture (20) and cancer cell lines (54), or studied chaperones that were co-regulated in cancer samples (37) or in aging (35), yet the fundamental structure of the functional network of chaperones in physiological human tissues was rarely analyzed. To identify core chaperones that were co-regulated, and thus likely functionally-related, in tissue contexts, we computed the pairwise correlation between the expression levels of each pair of chaperones across samples of the same tissue. We then constructed a functional network per tissue that contained interactions between highly-correlated core chaperones (r>0.7, Spearman correlation). Though core chaperons were uniformly expressed across tissues, their functional relationships changed markedly (Fig. 6A). For example, the two chaperonin subunits, CCT5 and CCT6A, were highly-correlated across all tissues (median correlation r=0.79), apart from testis where it was the lowest (r=0.54).

**Figure 6.**
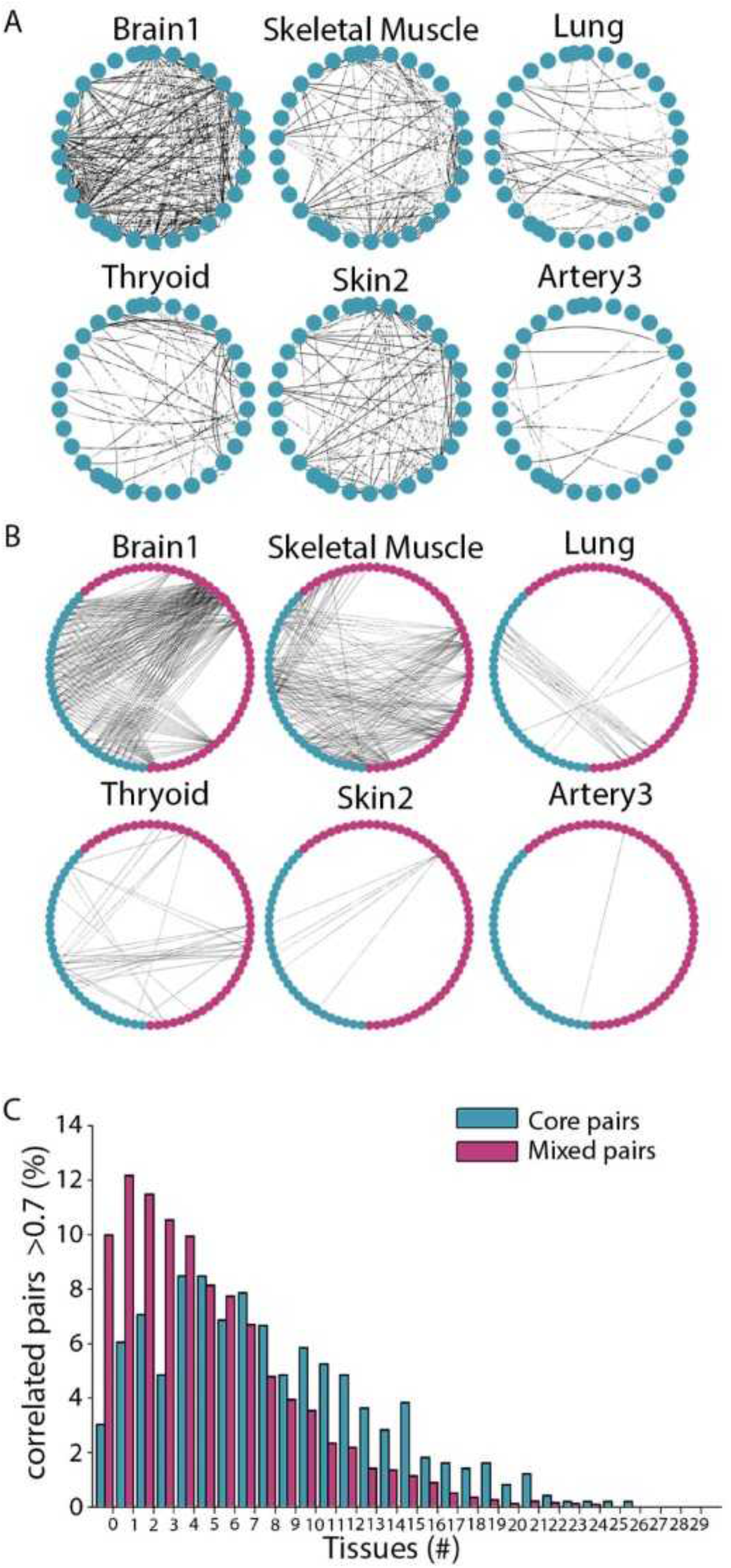
Chaperone functional networks differ between tissues. A. The functional networks of core chaperones in different tissues. Interactions connect chaperones with correlated expression levels in the respective tissue (r>0.7, Spearman correlation). Across tissues, core chaperones modulate their relationships with each other, thereby creating tissue-specific networks with distinct folding capacities. B. The functional networks of core and variable chaperones in different tissues. Only mixed interactions, which connect between core and variable chaperones, are shown. Core and variable chaperones are depicted as blue and pink nodes, respectively. Interactions connect chaperones with correlated expression levels in the respective tissue (r>0.7, Spearman correlation). Across tissues, core chaperones modulate their relationships with variable chaperones. C. The distribution of pairs of core chaperones (blue) and mixed pairs of core and variable chaperones (pink) by the number of tissues in which their expression levels were correlated (r>0.7, Spearman correlation). Pairs of core chaperones tend to be correlated across more tissues relative to mixed pairs (p=6.2E-5, Kolmogorov Smirnov test).

Next, we extended these functional networks to contain variable chaperones that were highly-correlated with core chaperones, which we refer to as ‘mixed’ relationships. The mixed functional networks were highly distinct from each other in terms of their highly-correlated pairs (Fig. 6B). For example, in testis, instead of being highly correlated with CCT6A, CCT5 was highly-correlated with its paralog, the variable chaperone CCT6B (r=0.89), pointing to a functional plasticity of the chaperone network. In agreement, high levels of CCT6A are highly correlated with a broad range of cancers, while CCT6B high levels are specifically correlated with testicular cancer (55). To quantify the tissue-specificity of each type of relationship at large-scale, we associated each chaperone pair with the number of tissues in which the pair was highly-correlated, distinguishing between core-core and mixed chaperone pairs. We found that the mixed relationships were significantly more tissue-specific than core-core relationships (p=6.2E-5, Kolmogorov-Smirnov test, Fig. 6C), giving rise to distinct, tissue-specific functional networks.

To facilitate interrogation of the functional relationships between chaperones across tissues, we created the ChaperoneNet webtool (http://netbio.bgu.ac.il/chapnet/). Users can query ChaperoneNet by chaperone and tissue, and obtain a graphical network representation of the functional relationships of the chaperone in that tissue. The network highlights core chaperones, marks chaperone families, and reports known disease associations.

## Discussion

Chaperones are basic components of all living organisms, and are thus commonly regarded as house-keeping genes (1). But, are all folding environments similar to each other? In a multicellular organism, owing to the specialized structures, functions and environments of the different cell types, folding requirements could differ greatly. This raises the question of whether the chaperone system is organized as a one size fits all, or whether the requirements of each tissue are met by a tailored system. To address this question, we set to systematically examine the expression landscape of molecular chaperones across physiological human tissues.

We first examined the expression of chaperones across tissues, as measured via RNA-sequencing by the GTEx consortium (30). We determined that most chaperones are expressed in all tissues, are highly expressed in most of them, and are more essential than other protein-coding genes, suggesting they act as consistent building blocks across all tissues (Fig. 1). Nevertheless, chaperones are also associated with tissue-specific phenotypes, since germline aberrations in ubiquitous chaperones or in ubiquitous aggregation-prone proteins can lead to tissue-specific pathologies, such as muscle disorders (Fig 2A) or neurodegeneration (48, 56–58). Indeed, we found that most chaperones were variably expressed across tissues, suggesting that the chaperone system is not simply common and consistent across tissues (Fig. 2B).

Although GTEx allowed high tissue-specific resolution analyses, this transcriptomic dataset is based on samples that were collected from recently-deceased donors and thus inherently limited, as the observed chaperone levels might represent stressful conditions. To test for this, we repeated analyses while excluding known stress-induced chaperones (Fig. S2). We identified similar, though somewhat weaker trends, suggesting that the impact of stress is partial. Additionally, transcript levels might not be indicative of protein levels, as the general correlation between them is not high, though is indicative, (59). Nonetheless, the high transcript levels of chaperones matched well with their high protein levels, which were previously measured in human cell lines (6). To further assess the relevance of the transcriptomic profiles, we profiled the proteomes of differentiating mouse myoblasts (Fig. 3A). Despite the difference in measured molecules (transcripts versus proteins) and species, we observed a strong agreement between the differential transcriptomic and proteomic profiles (Fig. 3B). Moreover, upregulated human chaperones were also enriched for *C. elegans* muscle chaperones (Fig. 3E), and for chaperones causal for muscle disorders (Fig. 2C).

A small subset of chaperones, which we denoted as core, did act as consistent building blocks across all tissues. Belonging to fundamental cellular processes, such as *de novo* folding maturation and assembly, ER-quality control and translocation across organelles, this core subset was significantly more essential for growth than other chaperones (Fig. 5B), and was 2.5-fold less likely to be causal for heritable disorders (9.6% versus 25.5%, Fig. 5C), and are therefore likely lethal when mutated. Notably, core and variable chaperones were not strongly associated with the classical organization of chaperones by family. Whereas main ATP-hydrolyzing chaperones were more consistent across tissues, most chaperones families had both core and variably expressed members (Fig. 4). Despite their consistent nature, core chaperones were involved in tissue-specific functional relationships both with other core chaperones and with variable chaperones (Fig. 6).

Altogether, our analyses expose a novel layer of functional organization: core chaperones, involved in basic cellular function shared by all cell-types, behave as a one size fits all, whereas variable chaperones respond to the tissue-specific demands and thus behave as a tailored system. We thus expand the previous functional view of chaperones, whereby the transition between prokaryotes and eukaryotes separated stress induced from constitutive chaperones (26). We propose that the transition to multi-cellularly further separated constitutive chaperones into core and variable chaperones, allowing for the modulation of the chaperone system in a tissue-dependent manner (Fig 7).

**Figure 7.**
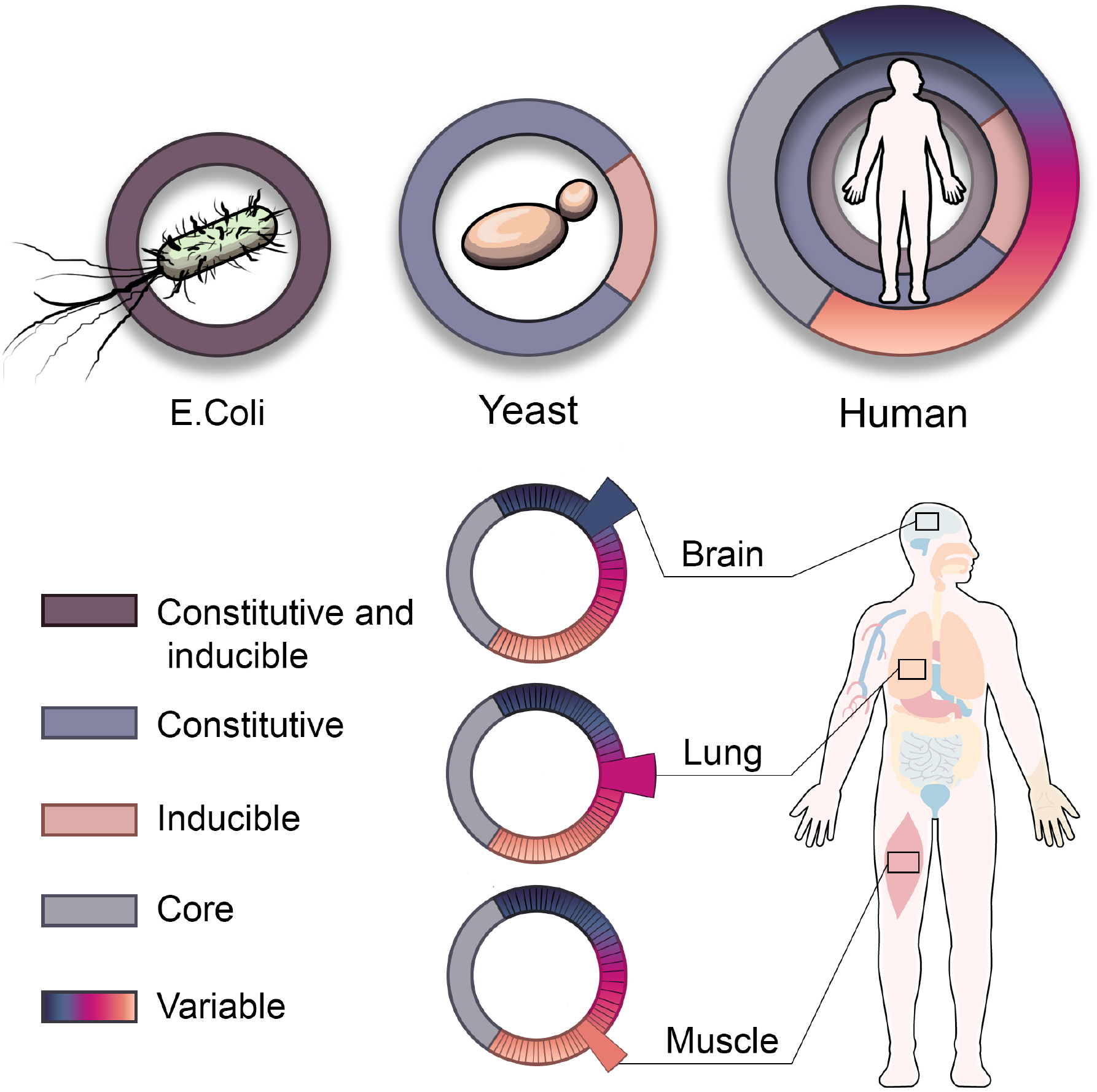
A layered architecture of chaperones across single- and multi-cellular organisms. The chaperone system of *E. coli* is composed of constitutive chaperones that can be induced upon stress (left). The chaperones system in yeast diverged to also include chaperones that are either constitutive or stress-inducible (middle). In multi-cellular organisms such as human, the chaperone system diverged further to also include core chaperones that are stable across tissues and variable chaperones that are differentially expressed across tissues (right), thereby giving rise to tissue-specific expression patterns, for example in brain, lung, or muscle.

The functional organization of chaperones is likely associated with their transcriptional regulation across tissue. In unicellular eukaryotes, chaperones required for *de novo* folding, such as CCT or prefoldin subunits and ribosome associated HSP70, were transcriptionally and functionally linked to the translational machinery (26), as might still be the case for certain core chaperones. Variable chaperones, in contrast, might be upregulated by tissue-specific transcription factors. In support, the main muscle differentiation transcription factor, MYOD1 (HLH-1 in *C. elegans*), was shown to drive the expression of chaperones in C2C12 mouse myoblast and *C. elegans* muscles (19, 60, 61).

Chaperones tissue-specific organization could offer an explanation to the tissue-specificity of chaperonopathies and protein-misfolding diseases (56, 58, 62). These diseases manifest clinically in few tissues, although most disease-causing genes and chaperones are expressed ubiquitously (Fig. 2A) (63, 64). In some cases, this could be due to the overexpression of causal genes in their disease-manifesting tissue (32, 65–67), as we observed for muscle diseases (Fig. 2C). We propose that it could also stem from chaperones collaborating or competing with each other over substrates, thereby leading to a proteostatic environment that is limiting to other proteins (68–72). Since both chaperones and substrates levels are altered between tissues, and since these alterations are functional, the proteostatic environment could both contribute to tissue-specific functionality and elicit tissue-specific phenotypes when perturbed, thereby giving rise to tissue-specific protein misfolding diseases.

Targeting ubiquitous chaperones can give rise to non-specific effects and has therapeutic limits (73). The analyses that we provide can potentially direct our focus to ubiquitous chaperones that are more sensitive to cell-specific modulation of their expression and thus are potential therapeutic targets for cell-specific disease. Formulating and quantifying the function of the chaperone system at large scale and within single cells is therefore one of the main challenges in proteostasis research.

## Materials and Methods

### Curation of an integrated list of human chaperones

We compiled a curated list of 195 known human chaperone, co-chaperone and folding enzyme genes, collectively named chaperones (Table S1). For that, we compared two curated lists of human chaperones: (1) a list of 332 chaperones curated by considering their biochemical properties and protein domains (35); and (2) a list of 194 chaperones complied based on sequence homology to conserved canonical chaperones and supplemented by a list of known co-HSP90 chaperones (6, 74). These lists included 114 genes belonging to well-conserved chaperones families, including sHsp (10 genes), HSP60/HSP10 (15 genes), HSP70 (13 genes), HSP90 (4 genes), prefoldin (9 genes), and co-chaperones Hsp40 (49 genes) and NEFs (14 genes). 106 of these chaperones appeared in both lists (~93% overlap). Less agreement between the lists was observed for the co-Hsp90 family, where only 24 (19%) chaperones out of 127 putative co-HSP90 chaperones appeared in both lists (~19% overlap). This inconsistency was due to the lack of a conserved domain shared by all members of the co-HSP90 family, and uniquely by them. For example, many known co-HSP90 chaperones have a tetratricopeptide repeat (TPR) motif, but not all TPR-containing proteins are chaperones (75). We therefore included 34 co-HSP90 chaperones that were previously shown to interact or function with HSP90 (15, 76, 77), three of which did not appear in any list and 22 appeared in both lists (~71% overlap).

The remaining genes in the initial lists (6, 35, 74), denoted as ‘others’, included members such as ER chaperones, mitochondrial chaperones, AAA+ and folding enzymes. For each member, we curated the literature to determine whether it functions as a chaperone that transiently assists the folding, unfolding, assembly or disassembly of proteins and protein complexes. For example, we excluded eight nuclear cyclophilins that appeared in both lists, as these were recently shown to constitute an integral part of spliceosomal complexes and were renamed as spliceophilin (78). Finally, we excluded genes that were annotated as pseudogenes or as noncatalytic proteins. For example, of the six HSP90 family genes, we excluded two pseudogenes (HSP90AA2 and HSP90B2P). Of the 47 chaperones that we included as ‘others’, 30 appeared in both lists (~64% overlap). Out of the 195 chaperones, only PPIAL4A did not meet the expression threshold of 1 TPM in any tissue and was therefore omitted from further analyses.

### Additional chaperone subsets

We annotated the functionality of core chaperones by manual curation of their Uniprot entries (79) and the literature. Data of curated, muscle chaperones in *C. elegans* were obtained from (19). Data of orthologous chaperones in *C. elegans* were obtained via the OrthoList2 tool for comparative genomic analysis between *C. elegans* and humans (80), and each human chaperone were fitted with the *C. elegnas* gene(s) that was identified as orthologue by the maximal number of programs. Data of orthologous chaperones in mouse were obtained from Mouse Genome Informatics (MGI) website (81).

Genes that are causal for Mendelian diseases were obtained from OMIM (82), and limited to genes with a known genetic basis. Manually curated data of the tissues that clinically manifest a Mendelian disease were obtained from (64). We additionally mined the literature for chaperones that are causal for muscle disorders. Chaperones that are induced by heat shock were obtained from a metanalysis study of several microarray data sets of stress-inducible genes probed for 167 chaperone genes (74).

### Human gene expression dataset

Data of transcriptomic profiles of human tissues measured via RNA-sequencing were downloaded from the GTEx portal (version 7). We analyzed samples from donors with traumatic injury fitting with sudden death, and avoided other death causes to limit the effect of treatment and stress on chaperones expression. We considered only physiological tissues (i.e., no transformed cells) with ≥ 5 samples. Brain sub-tissues were collapsed into three main regions including brain basal ganglia, largely cortex, and brain other, according to (83). Altogether, we analyzed 488 transcriptomic profiles and 29 tissues (Table S3). Genes were mapped to their Ensembl gene identifiers using BioMart (84), and filtered to include only protein-coding genes.

### Gene expression analyses

To analyze the expression pattern of genes across tissues, we considered genes as expressed in a tissue if their median expression level was ≥ 1 TPM. Results pertaining to expression thresholds of 5 or 10 TPM are presented in Fig. S1. The differential expression of genes across tissues was calculated as in (43). Specifically, we applied the TMM method by the edgeR package (85) to raw counts, to obtain a similar library size per sample. Genes with ≤ 10 raw counts per sample across all samples were removed before normalization. In each sample, we transformed the normalized counts using the voom method (86). To compute the differential expression of genes in a given tissue, we compared all profiles of that tissue to a background set containing profiles of all other tissues. Differential expression was then calculated by using the Limma linear model (87). Core chaperones were defined as chaperones that across all tissues had less than a 2-fold change in their expression; all other chaperones were considered variable.

### Essentiality analyses

The essentiality score of a gene was measured by the impact of its inactivation on growth rate relative to control, which was measured in four human cell lines (38). Data were available for all chaperones, except for DNAJB3, HSPA7 and HSPB9, and for 17,973 other protein-coding genes.

### Statistical analyses

To test the null hypothesis that the expression levels across tissues of chaperones and protein-coding genes were drawn from the same distribution we used the Kolmogorov-Smirnov test. We used the same test to compare between the essentiality scores of chaperones and protein-coding genes. To test the null hypothesis that gene expression levels of chaperones and protein-coding genes are similar, we used the Mann–Whitney U test. To test the null hypothesis that the essentiality scores of chaperones were similar to those of other protein-coding genes we considered per gene its essentiality scores in four human cell lines and applied the Kolmogorov-Smirnov test. To compute the statistical significance of the overlap between different gene sets we used Fisher exact test. Upon analyzing orthologous chaperones and the significance of the overlap, we considered all human chaperones in our data that had an orthologue in the respective species. To assess the correlation between the differential expression of human genes and mouse proteins in muscle samples we computed their Spearman correlation.

### Co-expression analyses

For each pair of chaperones and each tissue, we calculated the Spearman correlation between the expression levels of the two chaperones across all samples of that tissue. The functional network of a given tissue was set to include all pairs with correlation coefficient value ≥ 0.7. Networks were visualized via Cytoscape 3.5.1.

### ChaperoneNet implementation

The ChaperoneNet server was implemented in Python by using the Flask framework with data stored on a MySQL database. The website client was developed using the ReactJS framework and designed with Semantic-UI. The network view is displayed by the cytoscape.js plugin [ref Cytoscape.js]. The website supports all major browsers. Recommended viewing resolution is 1440 × 900 and above.

### Cell cultivation of C2C12 cells and Mass-Spectrometric Analysis

C2C12 cells (ATCC CRL1772) were grown in Dulbecco’s modified Eagle medium (DMEM) containing 10% Fetal Bovine Serum (FBS), Penicillin/Streptomycin (0.1 mg/ml) and L-glutamine (2 mM). For differentiation, cells were grown to 95% confluency and medium was changed to differentiation medium, containing 4% Donor horse serum (DHS) instead of FBS. During differentiation, the medium was changed at least every other day. Cells were harvested after 8 days, when they reached maximal differentiation. To separate myotubes from reserve cells, we used the property of myotubes to detach from cell culture plates under milder conditions than reserve cells. Plates were washed with PBS and incubated with trypsin diluted in DMEM medium without additives until most myotubes started detaching. Myotubes were then washed off the plate using PBS buffer. The remaining reserve cells were detached using 0.25% Trypsin and 0.05% EDTA solution. Both myotubes and reserve cell fractions were washed twice with 10 ml PBS and kept at −80°C. For Mass-Spectrometry, cells were suspended with 9M urea and 10 mM DTT solution and sonicated with a Cup-Horn sonicator for 2 min with 10 sec on and off cycles. After centrifugation to remove cell debris, samples were digested by trypsin and analyzed by LC-MS/MS on Q Exactive plus (Thermo). The data were analyzed using MaxQuant 1.5.2.8 versus *Mus musculus* part of the Uniprot database (Technion, Israel), and known contaminants were removed. Only proteins that were identified with at least 2 peptides were tested for significant differences. In the analysis of the correlation between mouse and human, we used only mouse proteins whose expression was measured at day 0 and day 8 and that showed significant differences, and their human homologs.

## Acknowledgements

This study was funded by Israel Ministry of Science and Technology grant 3-14337 to A.B.Z, S.C and E.Y.L.

## Author contributions

Conceptualization, N.S, A.B.Z and E.Y.L; Methodology, N.S, J.J, E.S, and R.B; Investigation, N.S, J.J, M.A, S.D, E.S, O.B and I.H; Formal Analysis, N.S, J.J, E.S, O.B and I.H; Visualization, E.V; Writing – Original Draft, N.S, A.B.Z and E.Y.L; Writing – Review & Editing, S.C, A.B.Z and E.Y.L; Supervision, A.B.Z and E.Y.L; Funding Acquisition, A.B.Z, S.C., and E.Y.L.

---

**Figure S1.**
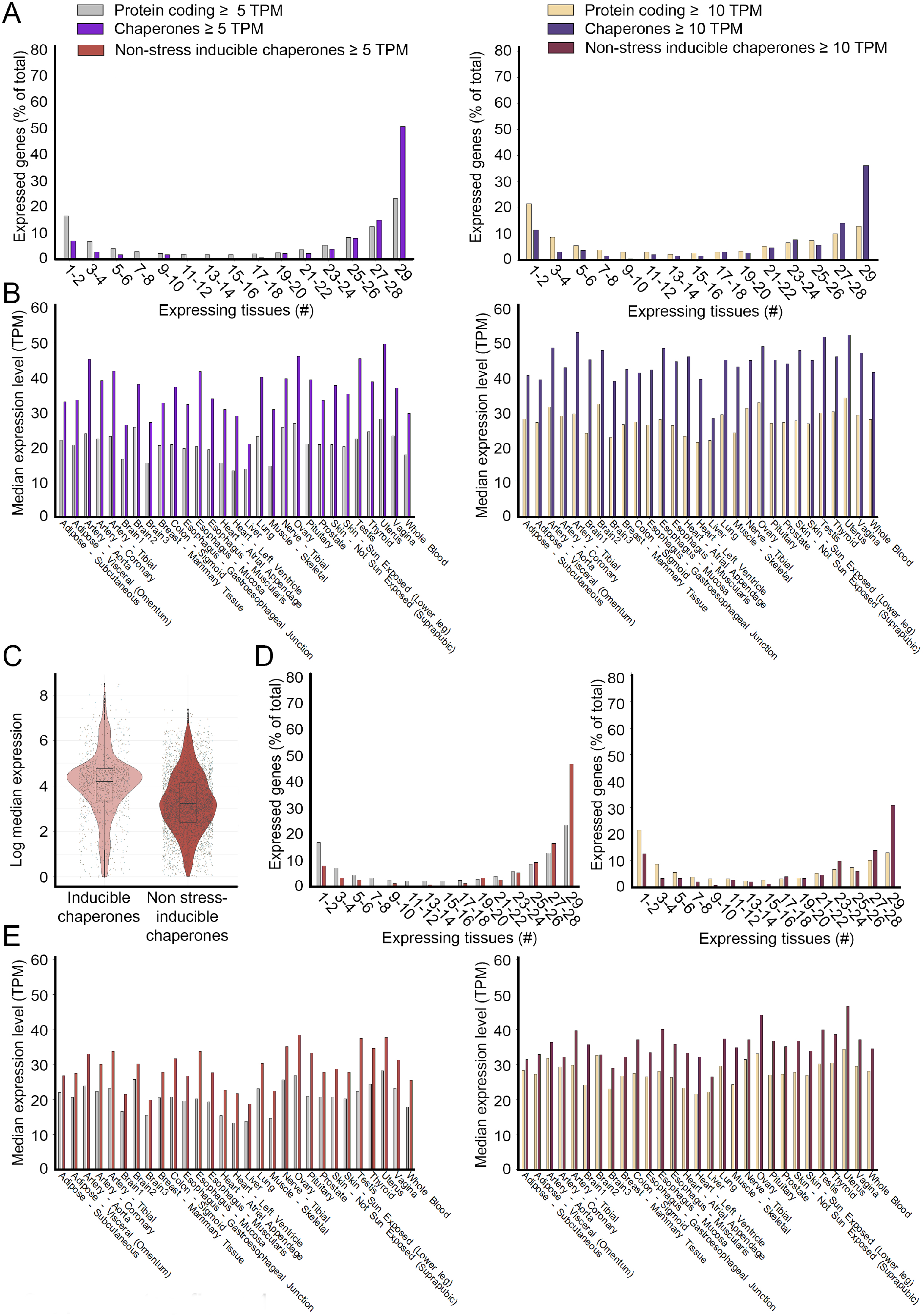
Chaperones are highly and ubiquitously expressed across tissues. A. The distribution of chaperones and other protein-coding genes by the number of tissues expressing them at a level ≥ 5TPM (left, 192 chaperones and 16,390 protein-coding genes) or ≥ 10TPM (right, 191 chaperones and 15,178 protein-coding genes). Chaperones are significantly more ubiquitously expressed than other protein-coding genes (p=6.7E-17 for both thresholds, Kolmogorov-Smirnov test). B. The median expression levels per tissue of chaperones and other protein-coding genes. Only genes expressed at a level ≥ 5TPM (left) or ≥ 10TPM were considered. Chaperones tend to be significantly more highly-expressed across all 29 tissues (p<3.45E-5, left, and p<0.0002, right, Mann Whitney test). C. The median expression levels of inducible versus non-inducible chaperones. Inducible chaperones are significantly more highly expressed (p<2.2E-16, Mann Whitney test). D. The distribution of non-inducible chaperones and other protein-coding genes by the number of tissues expressing them at a level ≥ 5TPM (left, 153 chaperones and 16,390 protein-coding genes) or ≥ 10TPM (right, 152 chaperones and 15,178 protein-coding genes). Non-inducible chaperones are significantly more ubiquitously expressed than other protein-coding genes (p=6.7E-17 for both thresholds, Kolmogorov-Smirnov test). E. The median expression levels per tissue of non-inducible chaperones and other protein-coding genes. Only genes expressed at a level ≥ 5TPM or ≥ 10TPM were considered. Non-inducible chaperones tend to be more highly expressed across tissues (p<0.02, left, or p<0.12, right, Mann Whitney test).

**Figure S2.**
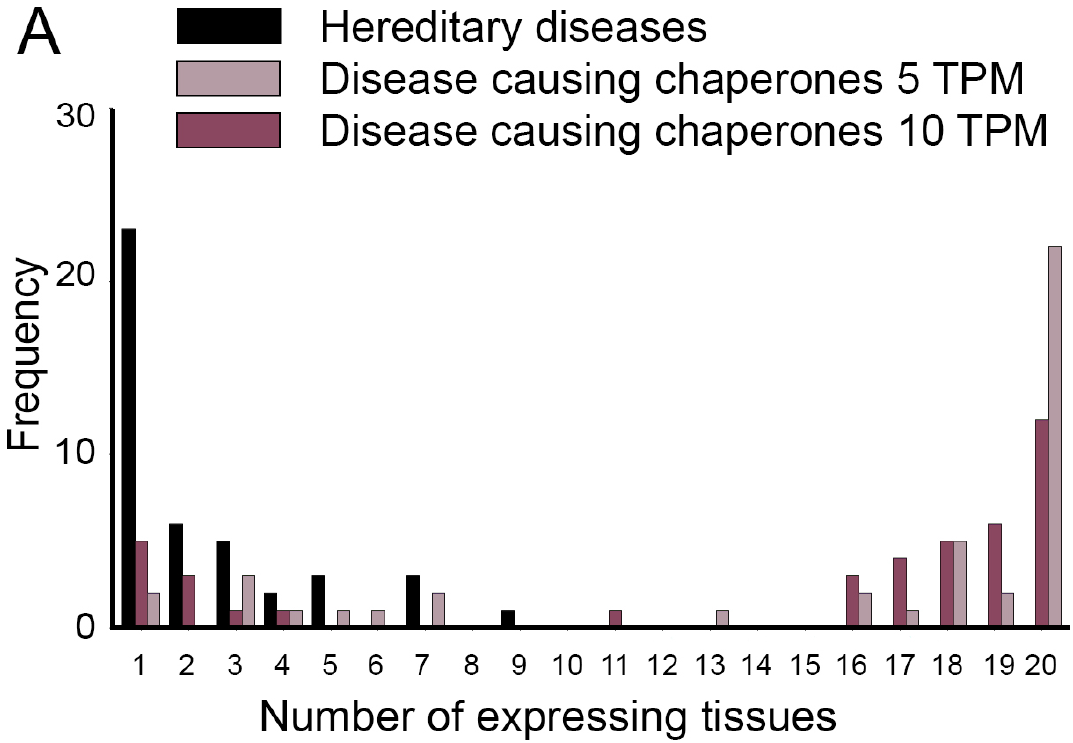
The distribution of chaperones that are causal for Mendelian diseases and the number of tissues in which their diseases manifest clinically. Most diseases are highly tissue-specific. In contrast, disease-causing chaperones are expressed ubiquitously across tissues upon considering expression thresholds of 5TPM or 10TPM. We united sub-parts of the same tissue (e.g., adipose subcutaneous and adipose visceral omentum were united into a single adipose tissue), therefore ending up with 20 united tissues.

